# Colonic spermidine promotes proliferation and migration of intestinal epithelial cells

**DOI:** 10.1101/2023.09.26.559404

**Authors:** Madison Flory, Angela Gao, Morgan Morrow, Ashfaqul Alam

## Abstract

The gut microbiome, comprising trillions of diverse microorganisms, profoundly influences the efficient development and maintenance of the intestinal barrier. While shifts in microbial composition are observed in Inflammatory Bowel Disease (IBD) and Colorectal Cancer (CRC), the causal relationship between these changes and resolution of inflammation remains elusive. Notably, IBD is not only marked by shifts in microbial composition but also by changes in microbial metabolites. Polyamines, produced by both gut bacteria and human cells, have emerged as potential regulators of gut pathology, cancer and mucosal repair. Investigating how elevated polyamine levels influence intestinal epithelial cells (IEC) can provide insights into their role in the regeneration of mucosal epithelia and restoration of gut barrier functions. To dissect the complex interplay between the gut microbiome, polyamines, and IEC, we focused on the overrepresented bacterium *B. uniformis* and its primary metabolite, spermidine. Here, we show that *B. uniformis*, a dominant member of gut microbiota, expands during the repair & resolution phase of the chemically induced chronic murine colitis. Furthermore, we found that the abundance of colonic polyamines was also altered, with spermidine being the abundant polyamine. Our RNA sequencing and transcriptomic analysis of cultured colonic epithelial cells demonstrate that spermidine regulates the expression of genes and pathways involved in different cellular functions, including proliferation, differentiation, adhesion, lipid metabolism, migration, chemotaxis, and receptor expression. We also found that spermidine stimulates the proliferation of cultured colonic epithelial cells in vitro. Additionally, our findings indicate that spermidine enhances the migrations of enterocytes. Our study emphasizes the crucial functions of the gut microbiome and polyamines in governing the functions of intestinal epithelial cells. Thus, these microorganisms and their metabolic byproducts hold promise as prospective therapeutic agents.

## Introduction

Although there are many components leading to the development and progression of inflammatory bowel disease (IBD), an extremely important factor in these diseases is the gut microbiome^1-5^. The complexity and constant variation inherently present in the relationship between the microbiome and the intestine provides a unique and important area of investigation as we seek to more deeply understand the causes and effects of IBD and attempt to develop new and innovative ways to treat these devastating lifelong diseases^1-7^.

There are two primary diseases that make up the IBD family: ulcerative colitis (UC) and Crohn’s disease (CD). Although the two share similar symptomology, and thus grouping together as IBD, they each demonstrate some characteristic differences isolating them from one another. In UC, inflammation is limited to the colon, and specifically, the mucosal layer of the colon^2^. Inflamed areas can be discrete, located among healthy portions of the colon. On the other hand, in CD, inflammation can occur anywhere along the gastrointestinal tract, and can occur in deeper layers of the intestines. Both disease states result in a similar set of symptoms, including diarrhea, abdominal pain, rectal bleeding, weight loss, and fatigue. Additionally, extraintestinal symptoms including inflammation in other parts of the body can sometimes occur^8,9^.

IBD is marked by cycles. At times, the disease is quiet, and patients experience a remission of symptoms. At others, flares occur, leading to a recurrence of symptoms and intestinal damage. There are many ways that IBD flares can be controlled and managed, including steroids, anti-inflammatory drugs, and biologic treatments, including monoclonal antibodies, all designed to reduce the amount of inflammation present^8,9^. The more cumulative length of time a person spends in a period of active inflammation, the higher their risk of developing later complications, including colitis-associated colorectal cancer (CAC), becomes. This becomes a critical observation as IBD diagnoses are increasing worldwide, especially in pediatric population. This results in an increase in lifetime inflammation, and will thus result in an increase in colorectal cancer in the future. In order to reduce this inevitable shift, we need to investigate the causes and contributing factors to IBD. Several have already been identified, but it remains to be determined exactly how each of these contribute to the disease process and which can be harnessed to use to benefit IBD patients^7-10^.

The gut microbiome is sometimes referred to as ‘the invisible organ’ due to its necessity, degree of complexity, and wide array of effects on both surrounding and distant tissues and organs. There are over 1 trillion individual microbes representing thousands of species, including bacteria, viruses, and fungi^1-5^. This collection of diverse microbes tells the story of our lives, varying based on someone’s birth, physical location, diet, infections and their treatment, and more. Changes in the microbiome over an individual’s life can have outsized impacts on their health, as the microbiome is known to affect vitamin synthesis, food digestion, drug metabolism, the immune system, angiogenesis, bone density, and the nervous system. There are many ways that the gut microbiome can impact health, and it is rather obvious that the gut microbiome will have an effect on IBD due to the direct proximity of the gut microbiome to the gut^10^.

The exact relationships and mechanisms between many individual bacteria and the body has yet to be determined, and we aim to more fully understand these relationships. It is known that gut bacteria can have effects on inflammation, and many high-level associations and trends have been studied, but what remains unknown is the actions and contributions of many individual microbes and their metabolic products to IBD^7-15^. The four major phyla of the gut microbiome – Firmicutes, Bacteroides, Proteobacteria, and Actinobacteria – make up the majority of the gut microbiome, with additional representation from less abundant phyla. In IBD, decreases in the Firmicutes phyla and increases in the Bacteroides and Proteobacteria are commonly seen. However, it is currently unclear whether these changes are driving resolution of inflammation in the gut or simply co-occurring with them.

Not only can characteristic changes in the microbes themselves be observed in IBD, but changes in their downstream metabolites are observed as well^10-15^. These products can go on to effect their own changes on intestinal epithelial cells, and can thus affect IBD for better or for worse. Many of these metabolic products are also produced by the eukaryotic cells of the colon, albeit in much smaller amounts. When the colon is in a state of dysbiosis and certain bacteria are overrepresented, so to will their metabolites be overrepresented. Increases in metabolites can have similar effects to the dysbiosis itself, changing the ways that the intestinal cells respond to certain stimuli and affecting the outcome of IBD^14,15^.

One such metabolite that can be overrepresented is polyamines. Human cells produce polyamines, but typically not in excess^6,8,35,36^. Gut bacteria also produce polyamines, and there exists mechanisms for these bacterially-produced polyamines to act on the intestinal cells. There are several different types of polyamines, all derived from amino acids, including spermidine, spermine, putrescine, and cadaverine. Polyamines are known to have impacts on many cellular processes, including cell division, differentiation, and migration^37,36^. Polyamine-producing bacteria are known to be elevated in IBD, resulting in an abundance of polyamines in the intestinal lumen. By investigating what the effects of polyamines are on intestinal epithelial cells, we can determine if these excess polyamines are contributing to the damaging effects of IBD, or are, in fact, a helpful metabolite in this case.

In order to determine the effects of the microbiome and its metabolites on IBD, we identified an overrepresented bacterial species – *B. uniformis -* and one of its corresponding metabolites – polyamines, primarily spermidine^42,44,45^. Focusing on a single species allowed us to further understand and isolate the effects from this overrepresentation on the system in order to understand the outcomes of enhancement of this particular species on a deep level. We focused on several key processes of IBD, including cell proliferation and migration, searching to understand how these necessary processes for intestinal wound healing were affected by one seemingly insignificant change to the microbiome. Here, we show that *B. uniformis*, a dominant member of gut microbiota, expands during the repair & resolution phase of the chemically induced chronic murine colitis. Furthermore, we found that the abundance of colonic polyamines was also altered, with spermidine being the abundant polyamine. Our RNA sequencing and transcriptomic analysis of cultured colonic epithelial cells demonstrate that spermidine regulates the expression of genes and pathways involved in different cellular functions, including proliferation, differentiation, adhesion, lipid metabolism, migration, chemotaxis, and receptor expression. We also found that spermidine stimulates the proliferation of cultured colonic epithelial cells in vitro. Furthermore, our results reveal that spermidine promotes the migration of enterocytes.

## Results

### Alterations in gut microbiota and colonic polyamines during the recovery phase of the chronic colitis

We previously demonstrated that the microenvironment of injured murine gut alters a local pro-restitutive microbiota^1^. However, we do not completely understand how the alterations of microbiota impacts colonic metabolites in the intestine undergoing repair after chronic inflammation. To model chronic intestinal inflammation and repair, mice were given three cycles of low dose of DSS (3%) for 5 days followed by 2 weeks of water. We collected stools on day 60 (1-week post DSS-removal; Figure A). DNA was purified from the stools as well as from adjacent intact mucosa and local luminal contents. Bacterial 16S rRNA genes (V4 region) were PCR-amplified and the amplicons were sequenced by high-throughput sequencing (HTS) platform. At the phyla level, microbiota analysis revealed that the stool microbiota consisted of a lower abundance of Firmicutes and a higher abundance of Proteobacteria and Bacteroidetes compared to the microbiota of the control mice. In addition, we found significantly increased abundances of *Ruminococcus (genus), Fusobacteria (phylum), Escherichia (genus), Prevotella (genus), S. gallolyticus*, and *Bacteroides uniformis* (Figure 1A-C). The initial increase in Bacteroidetes and *B. uniformis* prompted a further investigation into enrichment of specific bacterial species in these models. *Bacteroides* species are known to be producers of polyamine metabolites, indicating that we would expect an increase of polyamines in our CRC and chronic colitis models due to the increase in *Bacteroides* species. This was confirmed using MS/MS, where spermidine was found to be increased nearly 28-fold over the control group (Figure 1D). In a murine model, we found that when previously germ-free mice were colonized with *B. uniformis*, spermidine was increased (Figure 1E), further implicating gut bacteria, specifically, gut bacteria known to be increased in IBD, in excess polyamine production. These consistently characteristic increases in bacterial groups and their metabolites across our murine studies led us to further investigate the impacts of polyamine metabolites on human intestinal epithelial cells in relation to cellular pathways important for roles in IBD wound healing.

**Figure 1.**
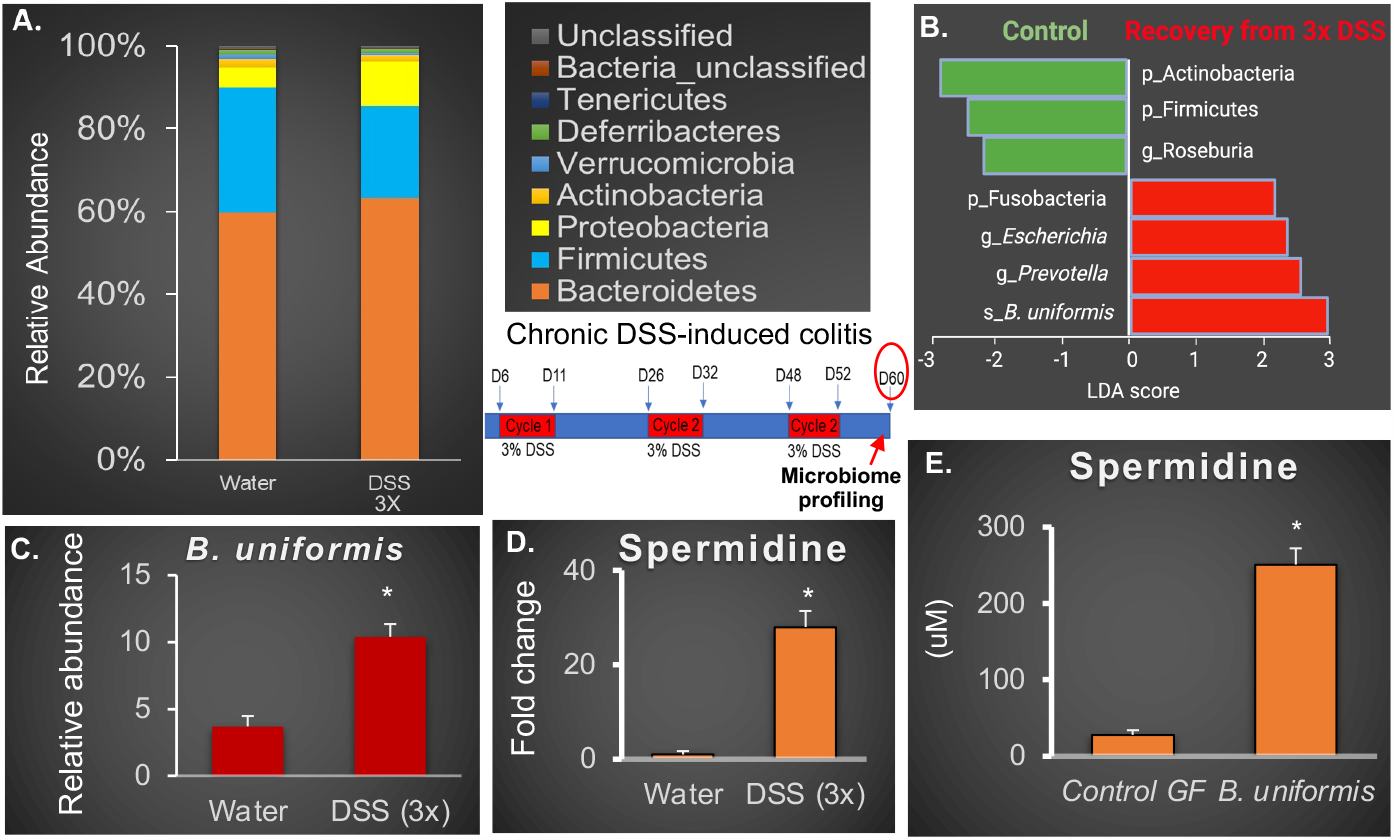
Microbiome profiling reveals characteristic changes in murine models of chemically induced colitis. (A) Mean relative abundance of bacterial phyla determined by 16s rRNA gene sequencing and microbiome analysis. (B) LDA scores reflect a positive association of bacterial taxa. (C) Specific increases of B. uniformis during the recovery phase post three cycles of DSS treatments. (D) Determination of spermidine in luminal contents of mice during recovery phase post three cycles of DSS treatments. (E) Determination of spermidine in luminal contents of ex-GF mice colonized with B. uniformis. N = 6-8 mice per group. ^*^P < 0.05; Student’s t-test.

### Spermidine alters the expression of genes & pathways critical for cell proliferation and migration of cultured intestinal epithelial cells

To evaluate the global impact of exogenous spermidine on the transcriptional response of cultured IECs, we performed unbiased RNA-sequencing using CaCO2 cells and measured the genome-wide mRNA expression profiles by quantitative deep sequencing of the RNA transcripts (Figure 2). We found marked differences in the mRNA profiles between spermidine-treated and vehicle-treated control cells. Clearly, numerous gene expression patterns in CaCO-2 cells were different from those in control. Among the differentially expressed genes, we observed in the increase in genes involved in IEC proliferation (Figure 2B). RNA sequencing of polyamine-treated Caco-2 cells revealed increases of many pathways and specific transcripts commonly implicated in mucosal repair and CRC, including increases in cellular proliferation, differentiation, and migration pathways, providing the impetus for investigation of these pathways in greater depth. As RNA sequencing had indicated polyamine treatment increased cellular proliferation pathways and transcripts, we set out to investigate this phenomenon at multiple levels. Using qRT-PCR to pair with our RNA sequencing, we investigated the changes in gene expression levels of several genes known to have roles in cell proliferation. The expression of CCND1 transcript was upregulated initially after 4 hours of treatment with spermidine, over a range of spermidine concentrations from 100 μM to 1 mM. The cyclin CCND1 helps to regulate the transition between the G1 and S phases of the cell cycle by forming complexes with CDK4 and CDK6, and is commonly upregulated during epithelial repair and in CRC. Together, this data indicates that exogenous polyamines are capable of increasing IEC genes and cellular signaling pathways involved in IEC proliferation.

**Figure 2.**
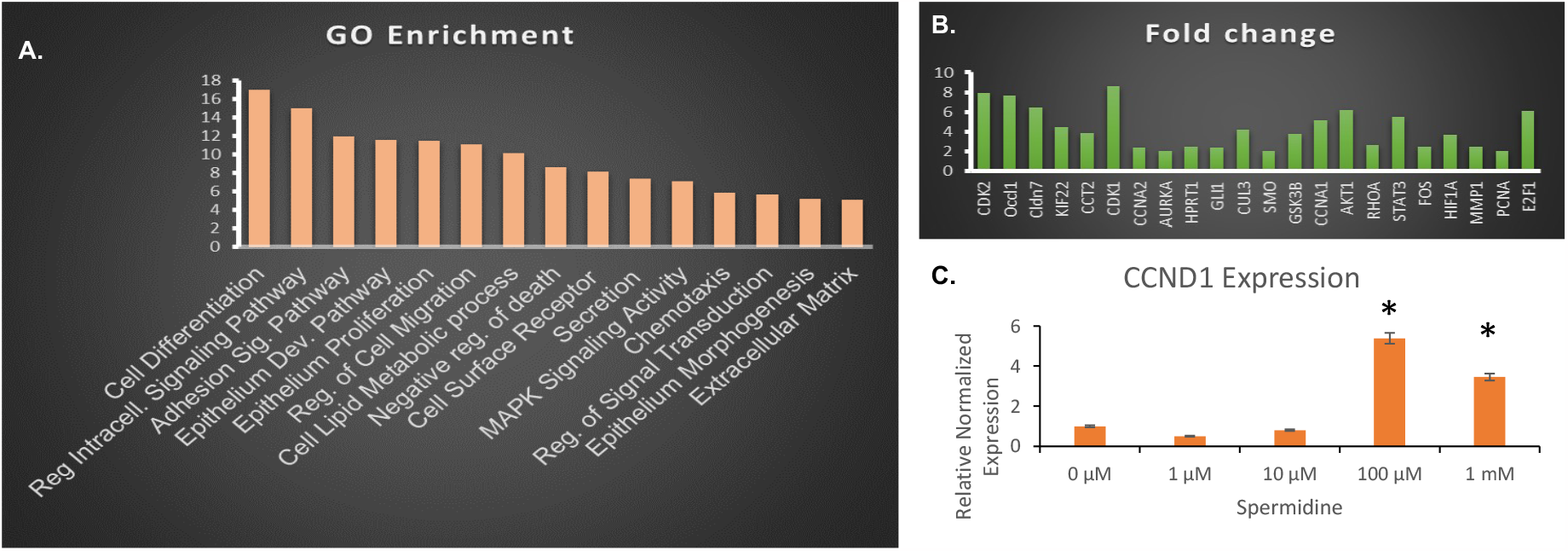
Spermidine alters the expression of genes & pathways critical for cell proliferation and migration of cultured intestinal epithelial cells. (A) GO Enrichment analysis of cellular pathways significantly increased in cultured IECs treated with spermidine. (B) Changes of specific genes involved in IEC proliferation. (C) qRT-PCR analysis demonstrating the increase of Cyclin D1 transcript expression in response to spermidine treatment. ^*^P < 0.05

### Polyamines increase cellular proliferation

Next, we examined IEC proliferation in vitro by EdU incorporation assay (Figure 3). Edu is a thymidine analog Edu and was used to identify cells in the S phase of cell division. DAPI was used to mark cell nuclei. Our data demonstrated that not only were increases seen in proliferation-related gene expression, but increased proliferation due to exogenous polyamine addition was identified in the cells themselves. By calculating an Edu/DAPI ratio, we estimated the proportion of actively dividing cells in a sample and investigate whether spermidine and other polyamines increase cell division (Figure 3B). The EdU incorporation assay was done using different concentrations of spermidine treatment to IECs. We found that the most effective concentration in increasing IEC division was 100 μM of spermidine (Figure 3A). Together, this data indicates that exogenous polyamines are capable of increasing IEC proliferation.

**Figure 3.**
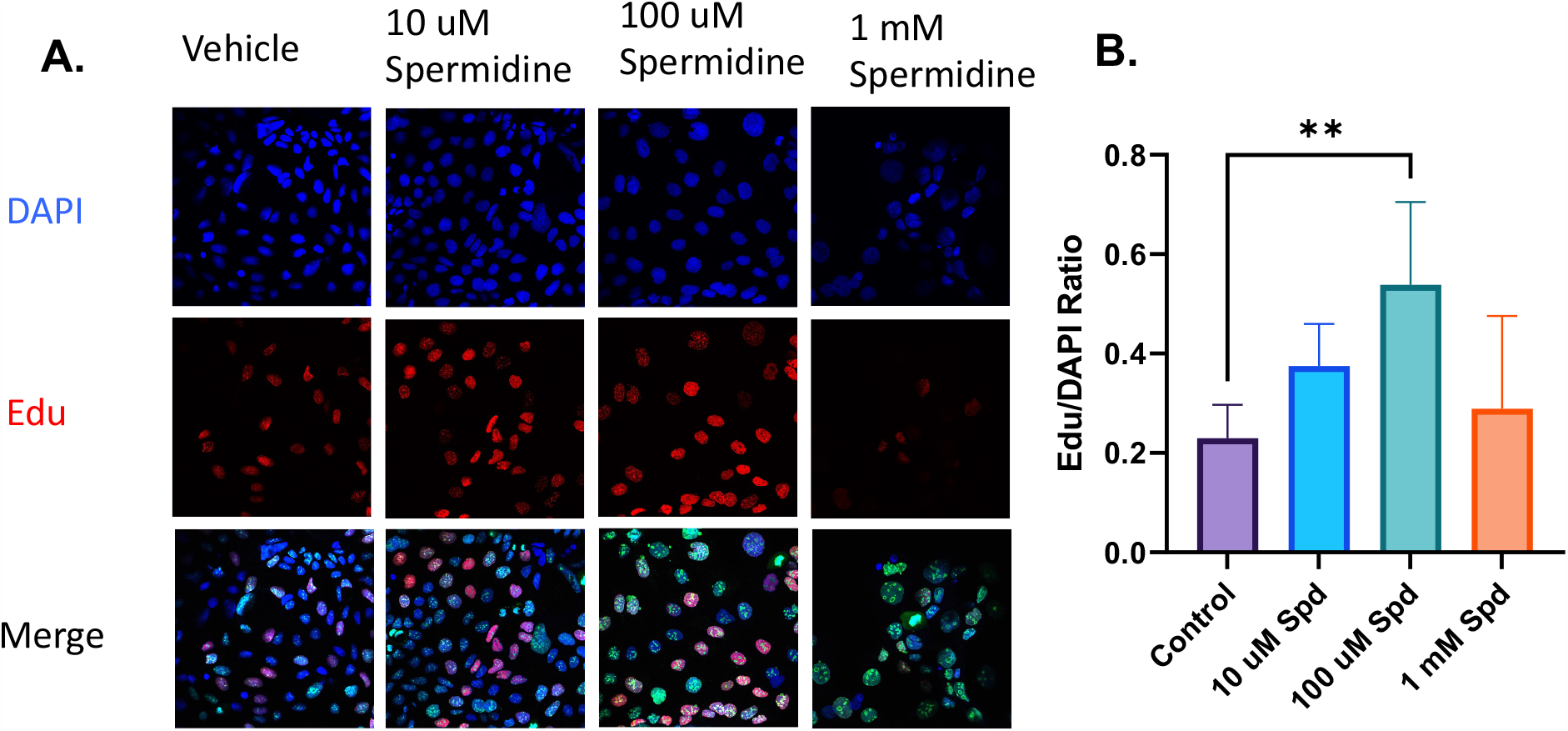
Spermidine stimulates the proliferation of cultured IECs. (A) Treatment with spermidine increases cellular proliferation as demonstrated by Edu incorporation assay. (B) Graph showing ratio of Edu /DAPI ratio. ^*^P < 0.05

### Polyamines increase cell migration

During restitution of colonic wounds, the epithelial cells migrate to repair the wound site. To determine if polyamines affected cellular migration, which is also a key hallmark of cancer primarily associated with metastasis, we performed a scratch assay (Figure 4A-B). This assay determines how well cells migrate in to fill a wound over time. On average, treatment with 100 μM spermidine increased the rate at which the cells recovered to fill the wound most effectively, but treatment with 1 mM spermidine also showed a significant increase at the 24-hour mark. We additionally investigated the activation of focal adhesion kinase (FAK) proteins involved in cell migration. We found that increased concentrations of spermidine stimulated phosphorylation of FAK (Figure 4C). Collectively, these data show that exogenous spermidine promote epithelial wound closure by stimulating phosphorylation of FAK.

**Figure 4.**
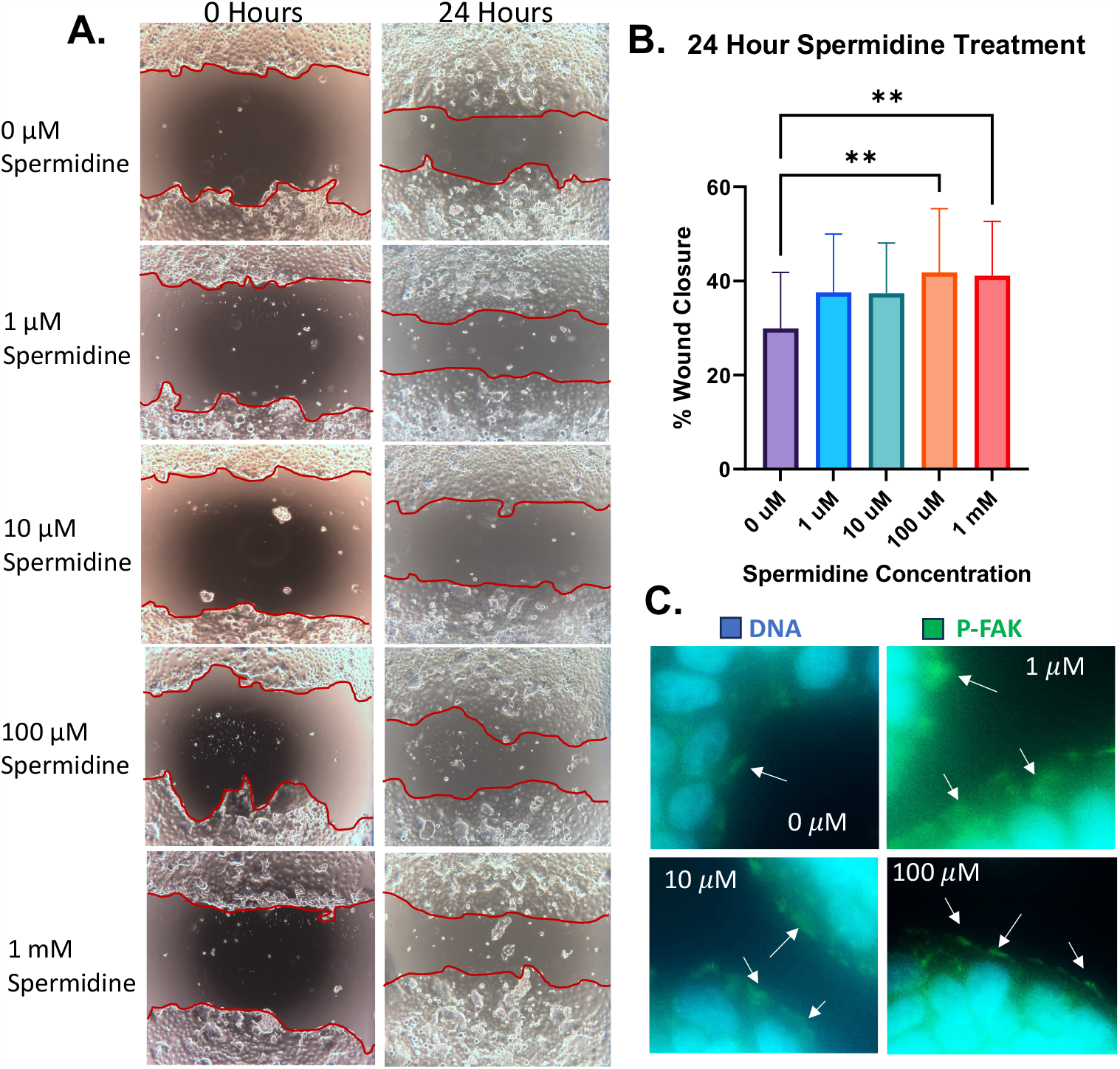
Spermidine promotes migration of IECs and increases phosphorylation of FAK. (A) CaCO2 monolayers were subjected to scratch wound assay in the presence of spermidine. Wound widths were determined at 0 and 24 hours. Photomicrograph shows representative results for control (Ctl) and spermidine – treated cells. (B) Graph shows percent wound closure. (C) Epifluorescence micrographs of IF staining highlighting phosphorylation of FAK in migrating CaCO2 cells with or without spermidine for 24 hours. ^*^P < 0.05

## Methods and materials

### Microbiome Study

DNA was extracted from stool samples (n = 5 mice per group) using a PowerSoil kit from MO BIO Laboratories (Carlsbad, CA)^1^. 16S rRNA genes were PCR-amplified from each sample using a composite forward primer and a reverse primer containing a unique 12-base barcode, designed using the Golay error-correcting scheme, which was used to tag PCR products from respective samples. We used primers for paired-end 16S community sequencing on the Illumina platform using bacteria/archaeal primer 515F/806R. Primers were specific for the V4 region of the 16S rRNA gene. The forward PCR primer sequence contained the sequence for the 5′ Illumina adapter, the forward primer pad, the forward primer linker, and the forward primer sequence. Each reverse PCR primer sequence contained the reverse complement of the 3′ Illumina adapter, the Golay barcode (each sequence contained a different barcode), the reverse primer pad, the reverse primer linker, and the reverse primer. Three independent PCR reactions were performed for each sample, combined, and purified with AMPure magnetic purification beads (Agencourt). The products were quantified, and a master DNA pool was generated from the purified products in equimolar ratios. The pooled products were sequenced using an Illumina MiSeq sequencing platform. Sequences were assigned to OTUs with UPARSE using 97% pairwise identity and were classified taxonomically using the RDP classifier retrained with Greengenes. After chimera removal, the average number of reads per sample was 19,717. A single representative sequence for each OTU was aligned using PyNAST, and a phylogenetic tree was then built using FastTree. The phylogenetic tree was used to compute the UniFrac distances. The PCoA analysis shown is unweighted.

### Metabolome Study

Stool or tissues from mouse distal colon were harvested and stored at -80°C. Then thawed samples were extracted by agitation with 1mL of degassed acetonitrile: isopropanol: water (3:3:2) at –20°C after which the soluble portion were recovered by centrifugation. Aliquots (450 μL) of that supernatant were used for each LCMS analyses. Specifically, we used an ultra-performance liquid chromatograph (Ultimate 3000) coupled to an Q Exactive HF, which incorporates an orbitrap analyzer. We injected on reverse phase as well as normal phase columns and collected in both positive and negative modes. In addition, we collected ion fragmentation (MS2) spectral to aid in feature identification. Raw mass spectrometry spectral data were collected for each biological replicate at each time point both in positive and negative MS modes. The data were processed using R to report integrated areas of each detected peak in individual samples and calculates the Welch’s t-test for two sample groups. For this study paired and unpaired non-parametric tests were carried out (Wilcoxon Rank Sum and Mann-Whitney). Features were listed in a feature list table and as an interactive cloud plot, containing their integrated intensities (extracted ionchromatographic peak areas) observed fold changes across the two sample groups, and p-values for each sample. Identifications were made by comparing retention time and tandem MS fragmentation patterns to the sample and a standard compound.

### RNA Extraction

Kit contents were prepared according to instructions. Cells were seeded at 5×10^4^ and grown overnight before treatment with water or spermidine for the appropriate amount of time. On ice, media was removed from each well and cells were washed with chilled D-PBS. 300 μL of RLT+ buffer was added and cells were scraped off of well using a pipette tip, then transferred into a microcentrifuge tube and vortexed for 45 seconds. 3 μL of β-mercaptoethanol was added to each sample before vortexing for an additional 45 seconds. Cell lysates were transferred to gDNA Eliminator columns and centrifuged at 10,000 rpm for 30 seconds. 600 μL of 70% EtOH was added to the flow-through and mixed by gentle pipetting. Sample was transferred to RNeasy spin columns and centrifuged for 15 seconds at 10,000 rpm, then 700 μL of Buffer RW1 was added to the columns and centrifuged at 10,000 rpm for 15 seconds. Flow-through was discarded and 500 μL of Buffer RPE was added before centrifuging at 10,000 rpm for 15 seconds. An additional 500 μL of Buffer RPE was added, this time centrifuging for 3 minutes at 10,000 rpm. The spin columns were placed into collection tubes, and 40 μL RNase-free water was added to the columns before centrifuging for 1 minute at 10,000 rpm. The flow-through was added back to the column and the centrifugation step repeated. Nanodrop was used to measure RNA concentration.

### qRT-PCR

Reverse transcription was performed according to the protocol in Bio-Rad’s iScript gDNA Clear cDNA Synthesis Kit. qRT-PCR was performed according to the protocol in Bio-Rad’s SsoAdvanced Universal SYBR Green Supermix following instructions for a 10 μL reaction. 2 μL of combined forward and reverse primers were used, along with 2 μL of nuclease-free H_2_O and 1 ng of cDNA template.

### RNA Sequencing

Cells were seeded in 24-well plates at 5×10^4^ and allowed to grow overnight before being treated with water or spermidine. RNA was extracted following protocol above. Quality of the RNA was evaluated using RNA 6000 Nano reagents on the 2100 bioanalyzer. RNA sequencing library preparation was performed utilizing the NEBNext Ultra RNA Library Prep Kit for Illumina by following the manufacturer’s recommendations (NEB). Sequencing libraries were validated on the Agilent 2100 Bioanalyzer System (Agilent Technologies) and quantified using Qubit 2.0 Fluorometer (Invitrogen) as well as by quantitative PCR (Applied Biosystems). The libraries were sequenced on an Illumina sequencer using a 2×150 Paired-End (PE) configuration. Raw sequence data (.bcl files) was converted into fastq files and de-multiplexed. Data were aligned and normalized using STAR aligner. Differentially expressed genes were identified by DEseq2. Gene pathway analysis was performed with the database for annotation, visualization, and integrated discovery (pathway enrichment analyses). We determined statistically significant differences in gene expression at the nominal level of significance (P < 0.05). We also evaluated the effect of Benjamini-Hochberg correction on raw p-values to account for multiple hypothesis testing.

### Edu Assay

ells were seeded at 1×10^5^ cells/mL and allowed to grow overnight before being treated with the appropriate concentration of water, spermidine, or DFMO for the appropriate length of time. Solutions were prepared according to kit protocol (Invitrogen C10337). Cells were incubated at 37° for 1 hour with 10 μM EdU. Cells were fixed using 4% paraformaldehyde for 15 minutes, then washed twice with 3% BSA. Cells were permeabilized with 0.5% Triton X-100 for 20 minutes and washed twice with 3% BSA. 0.5 mL per well of Click-It solution was prepared according to the protocol and added to each well, then incubated for 30 minutes at RT in the dark^41,1^. Cells were further stained using the IF staining protocol above beginning from the blocking stage.

### IF Staining

Cells were seeded at 5×10^4^ cells/mL and allowed to grow overnight before treatment with the applicable concentration of water, spermidine, DFMO, cisplatin, or acrolein for the appropriate amount of time. Cells were fixed in 4% paraformaldehyde for 15 minutes, then washed 3x with 1x PBS. Cells were incubated in blocking buffer at RT for 1 hour, then in primary antibody diluted in antibody dilution buffer overnight at 4°. Cells were washed 5x with 1x PBS, then incubated with secondary antibody diluted in antibody dilution buffer for 2 hours at RT in the dark. Cells were washed 5x with 1x PBS. Prolong Gold Antifade Reagent with DAPI was applied to each well, then a coverslip was applied. Antifade was allowed to cure overnight, then corners of coverslip were secured with nail polish. Confocal microscopy was used to image the cells.

### Scratch Assay

Cells were seeded at 1.2×10^5^ cells/mL and allowed to grow to confluency. A 200 μL pipette tip was used to scratch through the cells in a grid pattern. The wells were carefully washed with D-PBS and fresh media treated with the applicable concentrations of spermidine was added. Images of scratched locations were obtained immediately following the wounding process and every 6 hours after, continuing to 24 hours.

### Western Blot

Protein samples were combined with RIPA buffer to add equal amounts of protein to each well, then combined with loading buffer at a 3:1 ratio before boiling at 95°C for 5 minutes and centrifuging for 5 minutes at 12,000 g. Gels were run for 5 minutes at 50 V, followed by 35-40 minutes at 150 V, then transferred onto nitrocellulose membranes at 150 V for 35 minutes. Membranes were washed in 1xTBS for 5 minutes, then blocked in 1xTBST + 5% milk for 1 hour at RT. Membranes were washed 3 times in 1xTBST for 5 minutes, then incubated in primary antibody solution (1xTBST + 5% BSA) at 4°C overnight (p-FAK 1:2000, p-AKT 1:1000, Actin 1:2000). Membranes were washed 3 times in 1xTBST for 5 minutes, then incubated in secondary antibody solution (1xTBST + 5% Milk, 1 μL StrepTActin-HRP) for 1 hour at RT (Goat anti-rabbit 1:2000, goat anti-mouse (1:2000). Membranes were washed 3 times in 1xTBST for 5 minutes, before being incubated for 5 minutes in Clarity detection solution, then images were obtained. If stripping and reprobing was required, membranes were washed 3 times in 1xTBST for 5 minutes, then incubated with stripping buffer for 35 minutes at 50°C. Membranes were washed for 1 minute with tap water, then in 1xTBST three times for 5 minutes before repeating primary and secondary antibody dilution steps and obtaining images.

## Discussion

IBD is characterized by dysbiotic gut microbiota, compromised epithelial barrier function, chronic intestinal inflammation, and increased mucosal cytokines and immune cells infiltration. The symbiotic gut microbiota is intricately linked with intestinal health. Many gastrointestinal disorders, including IBD, are immensely impacted by the gut microbiota. However, the microbiota’s nearly 10 million genes and their metabolic functions remained largely unrecognized. Therefore, there is a critical vacuum of knowledge surrounding the specific but mechanistic roles of symbiotic microbial members, their metabolic functions, and the products they generate in the gut. Hence, there has been a remarkable interest in identifying the bacterial genes, pathways, and metabolic products, which may directly impact the functions of the intestinal epithelial cell (IEC) and immune cells of the gut.

The gut microbiota beneficially impacts intestinal homeostasis and, by extension, systemic organismal health. However, relatively little is known of how the host perceives non-pathogenic bacteria, or how the microbiota mechanistically influences gut biology by producing metabolic products.

Gut microbiota preferably catabolizes lysine, arginine, glycine, and the branched-chain amino acids (BCAA) to bioactive polyamines, including cadaverine, spermidine, and putrescine. Polyamines are primordial constituents of life, which are aliphatic amines synthesized by both eukaryotes and prokaryotes. Intracellularly synthesized polyamines are well characterized which regulate multiple cellular processes, including cellular proliferation, differentiation, and metabolic regulation. Polyamines are one of the major metabolites in the intestinal lumen.

Here, we describe a highly conserved metabolic pathway (the generation of polyamine) of gut microbiota that likely forms a fundamental component of the host–microbiota interaction. Timely and efficient intestinal mucosal wound repair, coinciding with resolution of inflammation, is critical in reestablishing the epithelial barrier and mucosal homeostasis. Resolution of inflammation and repair of the epithelial barrier are mediated by a delicate choreography of epithelium, immune cells and gut microbiota. Thus, an improved understanding of these biological processes are important in development of therapeutic strategies designed to promote recovery of epithelial injury in inflammatory states such as IBD. However, to date, little was known about the extent to which bacterial metabolites, specifically bacterial polyamine modulates IEC functions to control cellular proliferation and migration in the mammalian intestine. Here, we demonstrate how exogenous polyamine in the gut lumen modulates the epithelial proliferation and migration.

To dissect the complex interplay between the gut microbiome, polyamines, and IEC, we focused on the overrepresented bacterium *B. uniformis* and its primary metabolite, spermidine. Here, we show that *B. uniformis*, a dominant member of gut microbiota, expands during the repair & resolution phase of the chemically induced chronic murine colitis.

Furthermore, we found that the abundance of colonic polyamines was also altered during the resolution phase of the DSS-induced colitis, with spermidine being the abundant polyamine. Our RNA sequencing and transcriptomic analysis of cultured colonic epithelial cells demonstrate that spermidine regulates the expression of genes and pathways involved in different cellular functions, including proliferation, differentiation, adhesion, lipid metabolism, migration, chemotaxis, and receptor expression. We also found that spermidine stimulates the proliferation of cultured colonic epithelial cells in vitro.

Additionally, our findings indicate that spermidine enhances the migrations of enterocytes. Our study emphasizes the crucial functions of the gut microbiome and polyamines in governing the functions of intestinal epithelial cells. Thus, these microorganisms and their metabolic byproducts hold promise as prospective therapeutic agents.

